# Thermal Characterization and Preclinical Validation of an Accessible, Carbon Dioxide-Based Cryotherapy System

**DOI:** 10.1101/2024.03.01.582967

**Authors:** Yixin Hu, Naomi Gordon, Katherine Ogg, Dara L. Kraitchman, Nicholas J. Durr, Bailey Surtees

**Affiliations:** Kubanda Cryotherapy, Inc; Department of Radiology and Radiological Science, Johns Hopkins University; Department of Biomedical Engineering, Johns Hopkins University

**Keywords:** Breast Cancer, Cryotherapy, Global Health, Calorimetry, Thermal Analysis

## Abstract

To investigate the potential of an affordable cryotherapy device for accessible treatment of breast cancer, the performance of a novel carbon dioxide-based device was evaluated through both benchtop and in vivo canine models. This novel device was quantitatively compared to a commercial device that utilizes argon gas as the cryogen. The thermal behavior of each device was characterized through calorimetry and by measuring the temperature profiles of iceballs generated in tissue phantoms. A 45-minute treatment from the carbon dioxide device in a tissue phantom produced a 1.67 ± 0.06 cm diameter lethal isotherm that was equivalent to a 7-minute treatment from the commercial argon-based device which produced a 1.53 ± 0.15 cm diameter lethal isotherm. In vivo validation was performed with the carbon dioxide-based device in one spontaneously occurring canine mammary mass with two standard 10-minutes freezes. Following cryotherapy, this mass was surgically resected and analyzed for necrosis margins via histopathology. The histopathology margin of necrosis from the in vivo treatment with the carbon dioxide device at 14 days post cryoablation was 1.57 cm. While carbon dioxide gas has historically been considered an impractical cryogen due to its low working pressure and high boiling point, this study shows that carbon dioxide-based cryotherapy may be equivalent to conventional argon-based cryotherapy in the size of the ablation zone in a standard treatment time. The validation of the carbon dioxide device performed in this study is an important step towards bringing accessible breast cancer treatment to women in low-resource settings.

## 1. Introduction

Breast cancer is the most diagnosed cancer, with more than 2.3 million new cases each year, representing nearly a quarter of all cancer diagnoses in women [1]. Over 60% of breast cancer cases occur in low-and-middle-income countries (LMICs) and account for 70% of all breast cancer deaths [1]. Breast cancer has become one of the leading causes of morbidity and mortality in LMICs. In high-income countries, such as the U.K., U.S., Canada, and Northern Europe, women have an 85% five-year survival rate, compared to 50% in LMICs [2]. Major causes of this disparity are the geographic and socioeconomic constraints that limit access to standard healthcare facilities and treatments [3]. For many women, their socioeconomic status creates significant barriers to seeking proper care [2].

The current standard-of-care treatment for breast cancer in high-resource countries includes surgery, chemotherapy, and radiation. Unfortunately, these treatments are costly and often unavailable to women in low-resource settings [4]. Chemotherapy and radiation require expensive drugs, complex equipment, and robust supply chains. Surgical options, such as lumpectomy and mastectomy, require sterile operating rooms and surgical expertise that are not widely available in local clinics in LMICs. Globally, access to appropriate surgical care is alarmingly limited, with less than 25% of the population having access [5] and only 10% of LMICs having access to the 25 cancer medications on the WHO Model Lists of Essential Medicines [6]. Nearly 50% of African countries completely lack radiotherapy equipment [7]. In low-income countries, there is an average of one radiotherapy device per every 7 million people, compared to high-income countries where there is one per every 250,000 [8]. Aside from the restricted access to necessary resources for standard care, breast cancer patients in LMICs seeking treatment also face a significant financial burden, with average expenses of $1300 for surgery, $1400 for chemotherapy, and $1200 for radiation therapy [9].

Many LMICs are now promoting and adopting nationwide breast cancer screening programs [3,10], and, due to key patent expirations [11,12], affordable breast cancer diagnostics, such as handheld ultrasound and low-cost mammography, are now emerging [13-15]. However, despite advances in early detection, few affordable treatments are available to address the breast cancer burden in LMICs.

Cryotherapy is an established treatment method for many benign and malignant tumors and has been used since the 1970s in various cancer types, including prostate, kidney, liver, and lung cancer [16-17]. During the procedure, extreme cold is delivered via a probe to necrose cancerous tissue, and the body’s immune system subsequently breaks down the dead cells. Cryoablation is typically performed under local anesthesia/minimal sedation. Due to the small, minimally invasive cryoprobe and analgesic properties of cold, the procedure can be performed percutaneously under image guidance for deep organs and only leaves a scab at the treatment site. However, existing commercial cryotherapy systems are not feasible for use in LMICs due to their reliance on expensive and inaccessible cryogens such as liquid nitrogen and argon gas. These gasses necessitate specialized safe handling protocols, and the systems further require extensive specialized training to use the complex operating systems. Systems can cost upwards of $250,000 per unit [18] and more than $5,000 in single-use disposables per treatment [19].

This study investigates a carbon dioxide-based cryotherapy device by benchmarking its thermal performance against a commercially available argon-based system in both a calorimeter system and tissue phantoms. Additionally, an in vivo validation was conducted in a canine model to demonstrate proof-of-concept for future human studies with the device. This CO_2_ cryotherapy device meets many of the criteria for treating breast cancer in LMICs including affordability, ease of use, and operation without electricity. While CO_2_ is low-cost and abundantly available due to its use in the beverage industry, its use as a cryogen is considered limited because of its higher boiling point at -78.46°C as compared to more costly cryogens like liquid nitrogen and argon gas, which have boiling points of -195.8°C and -185.8°C, respectively. CO_2_ also has a lower working pressure of around 860 psi as compared to argon gas which is used at around 3000-4500 psi in cryotherapy systems [20]. Because the operating principle of most cryotherapy devices, the Joule-Thomson effect, relies on throttling a cryogen from high to low pressure, the lower working pressure of CO_2_ limits its cryogenic potential.

Low-cost cryotherapy based on cheaper cryogens such as gaseous and solid CO_2_ was explored during the early cryotherapy development of the mid-1900s, but interest in this technology decreased with the advent of liquid nitrogen- and argon-based systems [21]. Lower boiling point and higher-pressure cryogens allow for faster cooling, and as the per-minute operating room costs increased, shorter treatment times were prioritized for cost-effectiveness.

In prior work, the CO_2_ device underwent engineering optimizations, including the cryoprobe geometry and gas pre-cooling system, to fully harness the cooling power of gaseous CO_2_. Usability considerations were further implemented to make the device compatible with beverage- and food-grade CO_2_. The device’s efficacy in necrosing cancerous tissue and creating a clinically relevant sized iceball under heat load was validated in rat mammary tumors and swine liver, respectively [22].

In this study, the CO_2_-based device’s thermal performance was quantitatively compared to that of an argon-based cryotherapy system through bench tests, particularly the ability to reach similar freezing outcomes under heat load. First, a calorimeter test was performed for both devices to determine the total power output. Then, a bench test in tissue phantom simulating the human body’s thermal properties and heat load conditions was completed to evaluate the time efficiency in forming a clinically relevant sized iceball and to measure the lethal isotherm margins. Finally, one in vivo treatment using the CO_2_ device was completed in a client-owned dog with a spontaneously occurring mammary tumor to confirm the device’s ability to necrose cancerous tissue under physiological heat load.

## 2. Materials and Methods

### 2.1. Device Comparison

This study compares the thermal performance and cooling efficiency of a novel CO_2_-based cryotherapy device to a commercially available benchmark argon-based device indicated for use in human therapeutic applications. The metric used in this study to evaluate the ability of a cryotherapy device to induce cell death is the -20°C isotherm. The size of the -20°C isotherm generated by a cryotherapy device during treatment is widely accepted to correlate with the lethal zone in tissue [23-25]. Although CO_2_ cannot reach temperatures as low as argon due to its inherent thermal characteristics, it is still an effective cryogen as its boiling point is significantly below the lethal temperature of -20°C, making it an appropriate cryogen for many applications.

The CO_2_ device is entirely mechanical, consisting of a gas-regulating backend unit that directly connects to a standard pressurized CO_2_ tank. The device is operated by a simple on-off valve that controls the flow of gas to the system. High-pressure tubing connects the gas-regulating backend to a handle composed of two concentric tubes: a telescoping nozzle that throttles the incoming high-pressure CO_2_, and a larger pre-cooling chamber through which cooled, expanded gas is directed back to the backend and exhausted into the environment. An 8-gauge aluminum probe (Figure 1) is connected to the handle to conduct cold into the targeted treatment volume, generating an iceball and inducing tissue necrosis.

**Figure 1.**
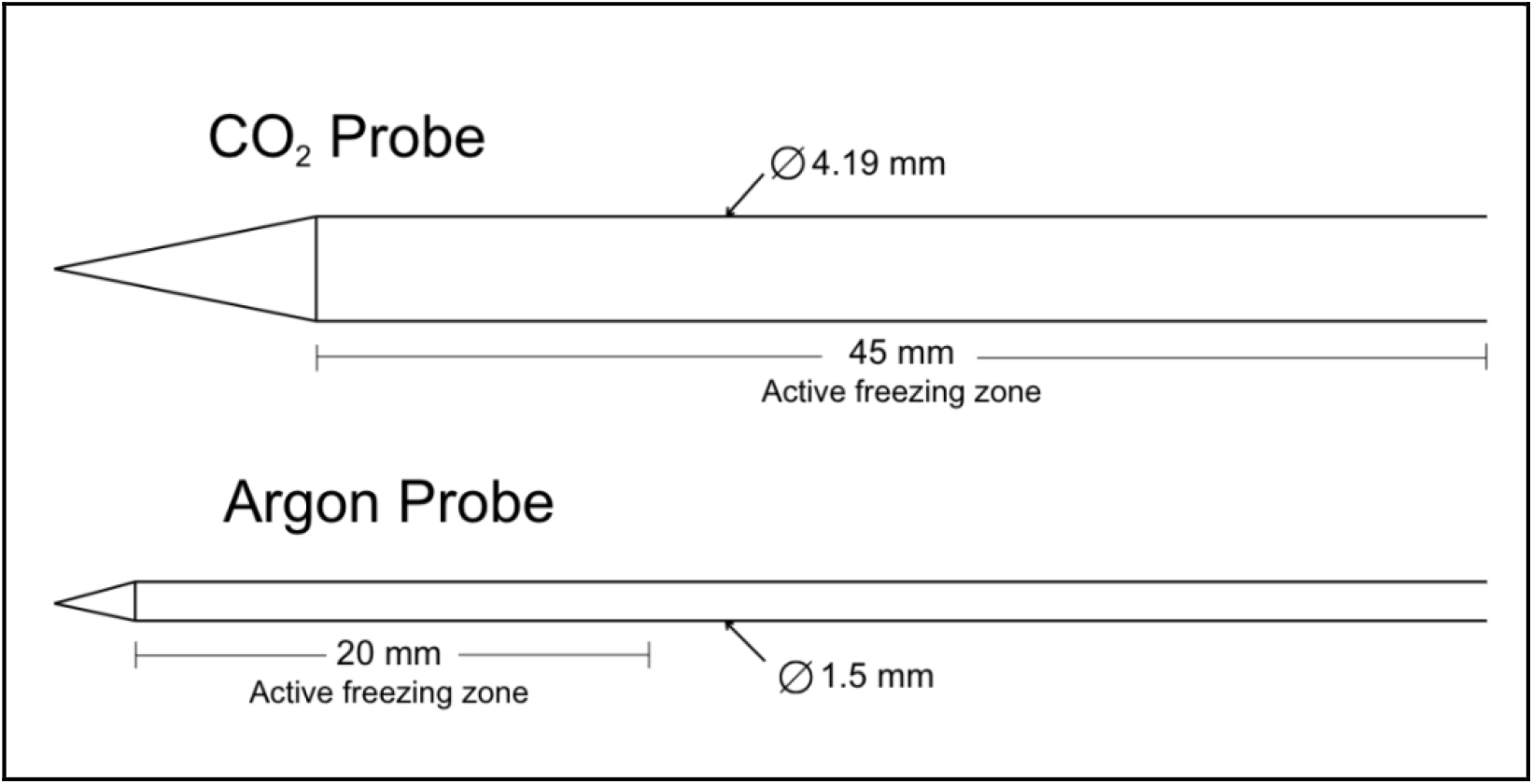
A comparison of the cryoprobe size, shape, and active freezing zone profiles is shown for the two devices. The CO_2_ device has a probe diameter of 4.19 mm, while the benchmark argon device has a probe diameter of 1.50 mm.

The benchmark argon device (Galil Medical, SeednetSystem, IceSeed probe) has a 17-gauge single-use stainless steel probe (Figure 1). Published performance specifications report its ability to generate a 2 cm diameter iceball with a -20°C 1.5 cm diameter isotherm, corresponding to the lethal zone during treatment. The benchmark device has an electronic control module for gas regulation and an active thaw feature that utilizes helium gas to warm the probe between freeze cycles [26]. Both devices have sharp-tipped probes designed to be inserted directly into the targeted tissue without the use of a scalpel.

Because of the difference in probe sizes of the two devices, methods of correction were introduced into each experiment to directly compare the experimental results of the two devices.

The primary focus of this study is to establish the efficacy of the CO_2_ device in comparison to a commercial device and to demonstrate that the CO_2_ device can induce clinically significant margins of necrosis in cancerous tissue.

### 2.2. Calorimeter Testing

The calorimeter setup consisted of two nested aluminum vessels, separated by insulating foam and sealed with an insulating lid. The inner cup was filled with 155 g of water at 65°C, and a stir bar was used continuously to ensure an even temperature distribution. To monitor the water temperature, a 40-gauge type-T thermocouple with PFA insulation (Omega Engineering) was fixed inside the internal vessel. Temperature data were recorded using a data logging thermometer (Omega Engineering, RDXL4SD). Before experimentation, the thermocouples were calibrated in accordance with manufacturer instructions against reference points of ice water and boiling water.

Three 140-second freezing trials were performed with each cryoprobe. During the trial, the cryoprobe was inserted into the calorimeter and allowed to freeze for 140 seconds. The temperature inside the calorimeter was recorded every 20 seconds. After each trial, the initial conditions were reset by reestablishing the water temperature to 65°C. The rate of change in temperature within the calorimeter was used to compute the cryoprobe’s cooling power, P, in Watts using the thermal energy equation,

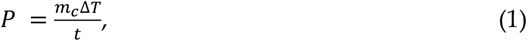

where *m* is the mass of the water in the calorimeter, *C*_*p*_ is the specific heat capacity of water (4.184 J/g°C), *ΔT* is the change in temperature, and *t* is the time interval. The cooling power measured from each of the three intervals was averaged to calculate the overall cooling power of the cryoprobe.

Cooling power over the surface area of the active freezing zone was normalized between the two systems to correct for the differences in probe size.

### 2.3. Tissue Phantom with Heat Load Testing

#### 2.3.1. Tissue Phantom

A benchtop tissue phantom was developed to simulate the thermal conditions of tissue during a cryotherapy treatment. A beaker containing 234 g of ultrasound gel was heated to 37°C and placed in a heated water bath that circulated at 37°C to simulate the heat load and perfusion conditions of a body with a constant core temperature (Figure 2).

**Figure 2.**
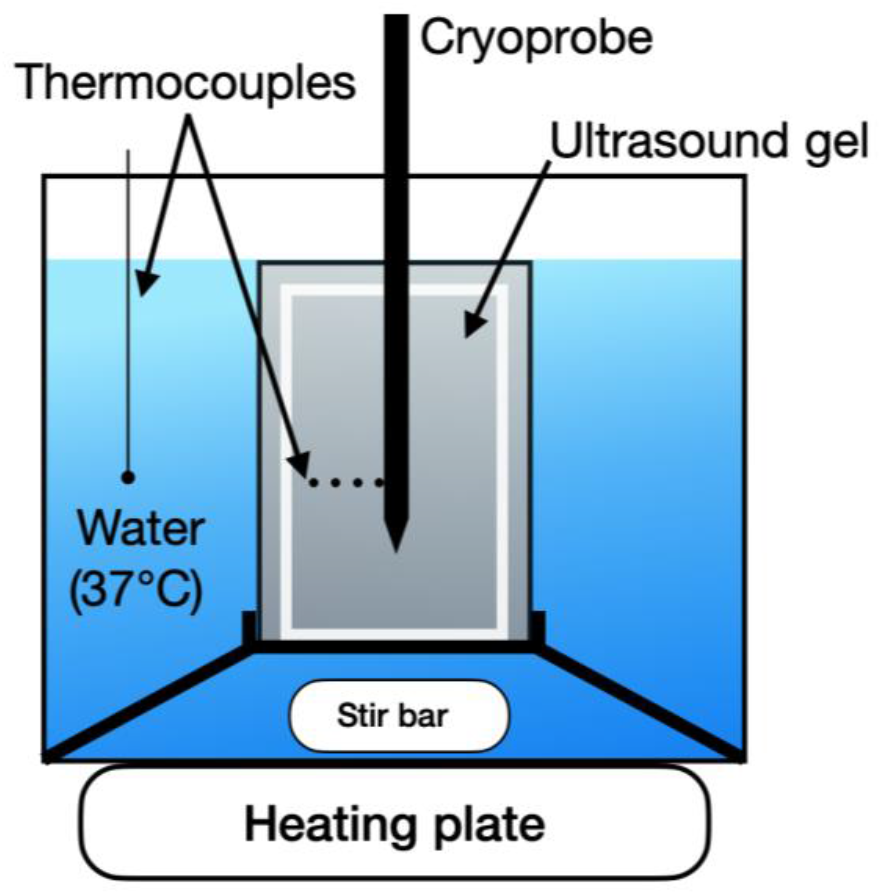
Temperature distribution testing setup. The cryoprobe is placed in a custom fixture containing heated ultrasound gel with four thermocouples at fixed radial distances from the probe surface. The ultrasound gel is surrounded by a 37°C water bath maintained with temperature monitoring and control, simulating the heat load conditions of a body.

A fixture centered the probe in the ultrasound gel and held the thermocouples at fixed radial intervals. To account for the difference in diameter between the CO_2_ probe (4.19 mm) and the argon probe (1.5 mm), a separate fixture was designed for each probe, and temperature measurements were taken at equivalent distances of 1 mm, 4 mm, 7 mm, and 10 mm from each probe’s outer surface (Figure 3), recorded with a 0.5 Hz sampling rate.

**Figure 3.**
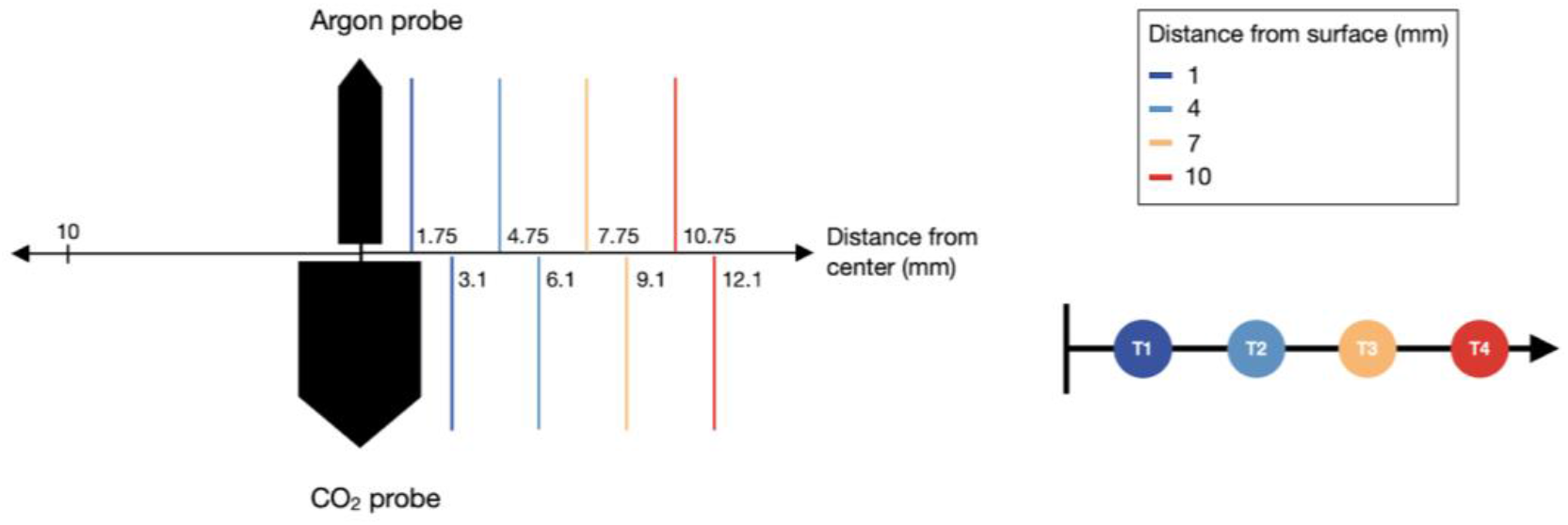
Positions of the four thermocouples. Left: A comparison of the center distance of the thermocouples for the argon and CO_2_ probes. Right: A diagram showing the distance of each thermocouple from the probe surface.

Preliminary tests were performed to determine the specific vertical position on each probe corresponding to the maximum diameter of the iceball. The thermocouple fixture was then aligned with this position during subsequent testing. Additionally, a fifth thermocouple was positioned 1 cm away from the probe’s center to monitor the time for the iceball to achieve a 2 cm diameter.

To measure an isotherm of clinically relevant size, the testing duration ranged from 7 minutes for the argon device to 45 minutes for the CO_2_ device. A total of 10 trials were conducted for each probe, and the temperature distributions were evaluated in Matlab. Minor fluctuations in the temperature data were smoothed with second-degree Savitzky-Golay filtering [27].

#### 2.3.2. Mathematical Modelling

The tissue phantom study aimed to determine the size of the -20°C isotherm for each device after a specified testing period. Because temperature values were recorded at discrete thermocouple locations, a curve-fitting model was used to approximate the temperature values across the entire iceball diameter perpendicular to the probe.

The tissue phantom was designed to simulate the thermal properties and heat load conditions of human tissue. To evaluate the theoretical temperature distribution in the tissue phantom, we began with the Pennes Bioheat Equation, which models the thermal dynamics within biological tissues:

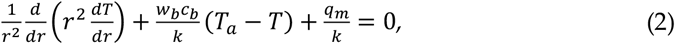

where *r* is the radius from an infinitesimal point source of cooling, *w*_*b*_ is the perfusion rate per unit volume of blood, *c*_*b*_ is the specific heat of blood, *k* is the thermal conductivity of tissue, *T* is the tissue temperature, Ta is the arterial temperature, and *q*_*m*_ is the metabolic heat generation per unit volume [28].

In the experimental tissue phantom, metabolic heat, arterial blood heating, and blood perfusion were not emulated, which reduces the bioheat equation to a simple conduction equation:

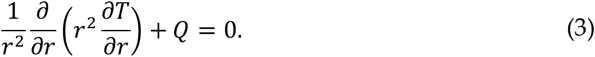

Solving this conduction equation in 1-dimensional cylindrical coordinates results in an equation of the form

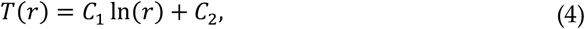

where *C*_*1*_ and *C*_*2*_ are constants [29].

The temperatures measured by each of the four thermocouples were averaged over 10 trials, and the temperature data at the end time points of the testing periods were evaluated as steady-state temperature distributions. A curve of the mathematical form in Equation 4 was fit to the four data points. The distance of the -20°C isotherm from the probe surface was determined from the best-fit curve.

To analyze the variation in temperature performance of the probes, the 95% confidence interval of the temperature data from each thermocouple was computed. Curves of the mathematical form in Equation 4 were fitted to the upper and lower confidence interval bounds and the points where each fitted curve intersected the -20°C isotherm were determined.

### 2.4. In Vivo Testing

#### 2.4.1. Cryotherapy Procedure in Canine Model with Spontaneously Occurring Mammary Cancer

The animal study protocol was approved by the Animal Care and Use Committee at Johns Hopkins University, and informed consent of the dog’s owner was obtained. The efficacy of the CO_2_ cryoablation device was validated in a 9-year-old 7.8 kg mixed-breed dog presenting with a ∽2.5 cm cystic mass on the right cranial mammary gland. A fine needle aspirate confirmed that the mass was an epithelial neoplasia with inflammation and cystic formation.

The dog was sedated with fentanyl (5 μg/kg IV) and received maropitant citrate (1 mg/kg IV) as an anti-nausea medication. The dog was then inducted with midazolam (0.25 mg/kg IV) followed by propofol (4 mg/kg IV). The dog was intubated, and general anesthesia was maintained with isoflurane with mechanical ventilation at ∽12-15 breaths per minute. Physiological monitoring during the procedure included temperature, blood pressure, ECG, oxygen saturation, and end-tidal carbon dioxide.

The dog was shaved, sterilely prepped, and draped, and the cryoprobe was inserted into the mass (Figure 4). Sterile saline was injected between the mass and skin to displace the skin as a preventative for cold damage to the skin. A small stab incision was made into the skin prior to the insertion of the CO_2_-based cryoablation probe. Following the recommended treatment dosage for a 3 cm mass given for both the CO_2_ device and the argon device, the cryoablation procedure was performed with two 10-minute freeze periods separated by a 7-minute passive thaw period. Throughout the procedure, the surgical site was observed for signs of cold damage to the skin, and the mass was manually palpated to monitor iceball growth. Ultrasound gel was placed on the skin and at the probe entry site, and room-temperature sterile saline was used to irrigate the skin to prevent thermal damage to healthy skin. After the probe was removed, the entry site was closed with a skin staple. The dog was allowed to recover and was returned to the owner when fully awake. Surgical resection of the mass was performed 14 days after cryoablation under general anesthesia.

**Figure 4.**
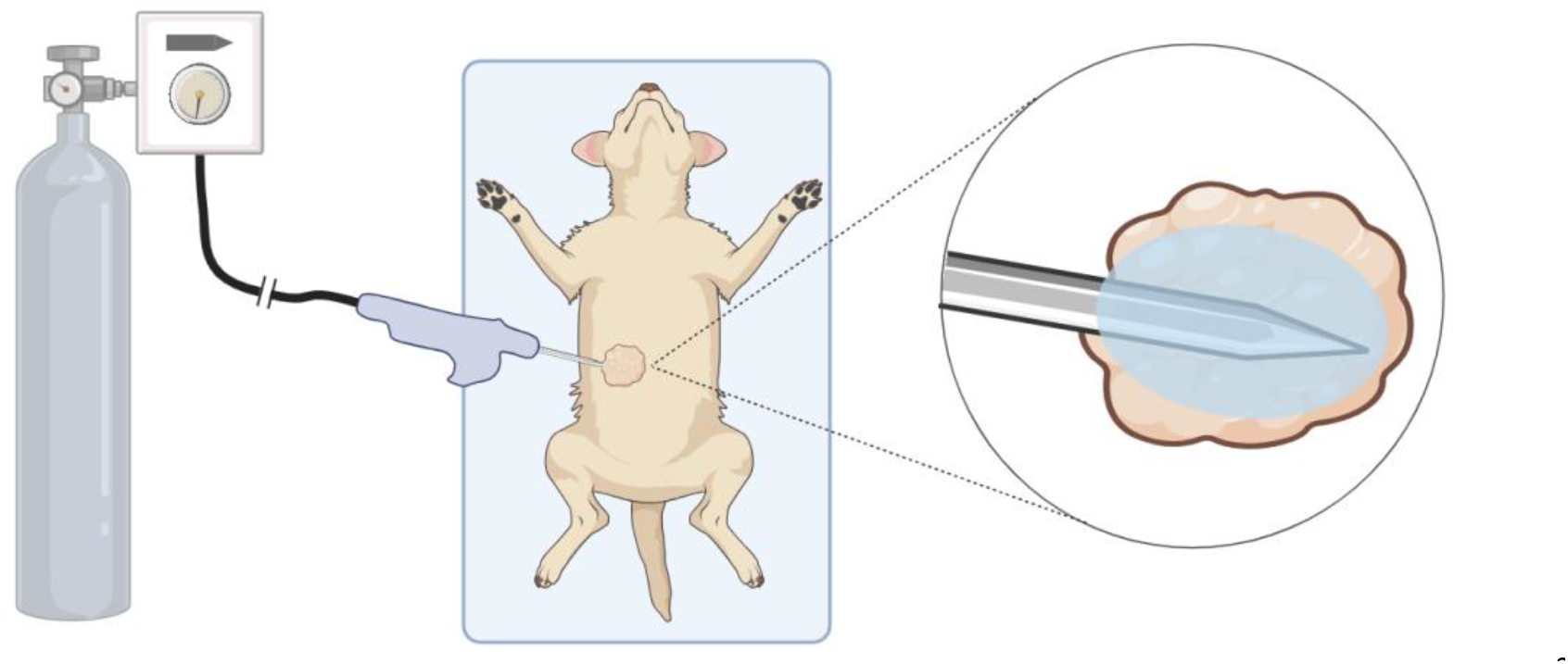
Diagram of the cryotherapy procedure setup. Left: the device is connected to a standard CO_2_ tank. Center: The subject is sterilely draped and prepped, and the probe is inserted into the target mass. Right: The device is turned on, and an iceball begins to grow around the probe tip inducing necrosis in the mass.

#### 2.4.2. Pathology Methods

The surgically resected tissue was fixed in 10% neutral buffered formalin and then sliced along the plane parallel to the path of probe insertion. Slides were made from paraffin-embedded tissue sections. The slides were stained with hematoxylin and eosin (H&E) and digitized for histopathological analysis. Tissue was identified as necrotic when either no viable nuclei were detected or the cellular architecture was entirely disrupted (Applied Pathology Systems).

## 3. Results

### 3.1. Calorimeter Results

The average cooling power of each device, as measured over 3 trials, is shown in Table 1. The calculated surface area of the effective freezing zone for each probe was used to normalize the cooling power between devices. This data demonstrates that the CO_2_ device generates a much greater overall cooling power. However, when normalized over the surface area of the effective freezing zone, the two cooling powers become more comparable. The argon device has a higher cooling power per unit surface area.

**Table 1.**
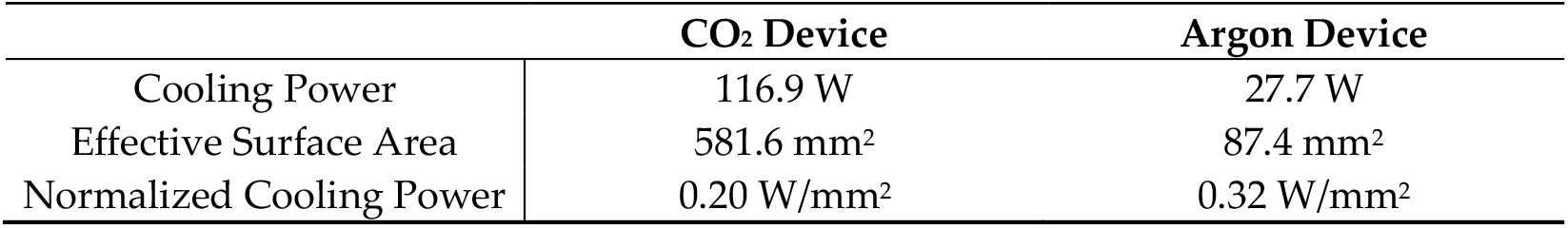
Comparison of normalized and overall cooling power between the CO_2_ device and the argon device.

|  | CO<sub>2</sub> Device | Argon Device |
| --- | --- | --- |
| Cooling Power | 116.9 W | 27.7 W |
| Effective Surface Area | 581.6 mm<sup>2</sup> | 87.4 mm<sup>2</sup> |
| Normalized Cooling Power | 0.20 W/mm<sup>2</sup> | 0.32 W/mm<sup>2</sup> |

### 3.2. Temperature distribution results

The temperature values measured by each of the four thermocouples at distance r from the probe surface are shown in Figure 5. The mean and 95% confidence intervals of the temperatures reached at each thermocouple at the end of the testing periods are shown in Table 2. The temperature at 7 mm from each probe surface (corresponding to an overall diameter of 18.2 mm and 15.5 mm for the CO_2_ and argon devices, respectively) reached similar temperature values at the end of the testing periods. For the CO_2_ device, the final average temperature at 7 mm from the probe surface was -20.09°C, while for the argon device, the temperature was -20.90°C. Figure 6 shows the iceballs generated by each device during testing.

**Table 2.** Mean temperature with 95% confidence intervals of each thermocouple at the end of the testing period for each device.

| Thermocouple Location | CO<sub>2</sub> (°C) | Argon (°C) |
| --- | --- | --- |
| 1 mm | -41.62 ± 1.31 | -78.19 ± 3.19 |
| 4 mm | -25.89 ± 0.60 | -46.76 ± 6.79 |
| 7 mm | -20.09 ± 0.57 | -20.90 ± 2.28 |
| 10 mm | -12.81 ± 0.58 | -1.30 ± 3.376 |

**Figure 5.**
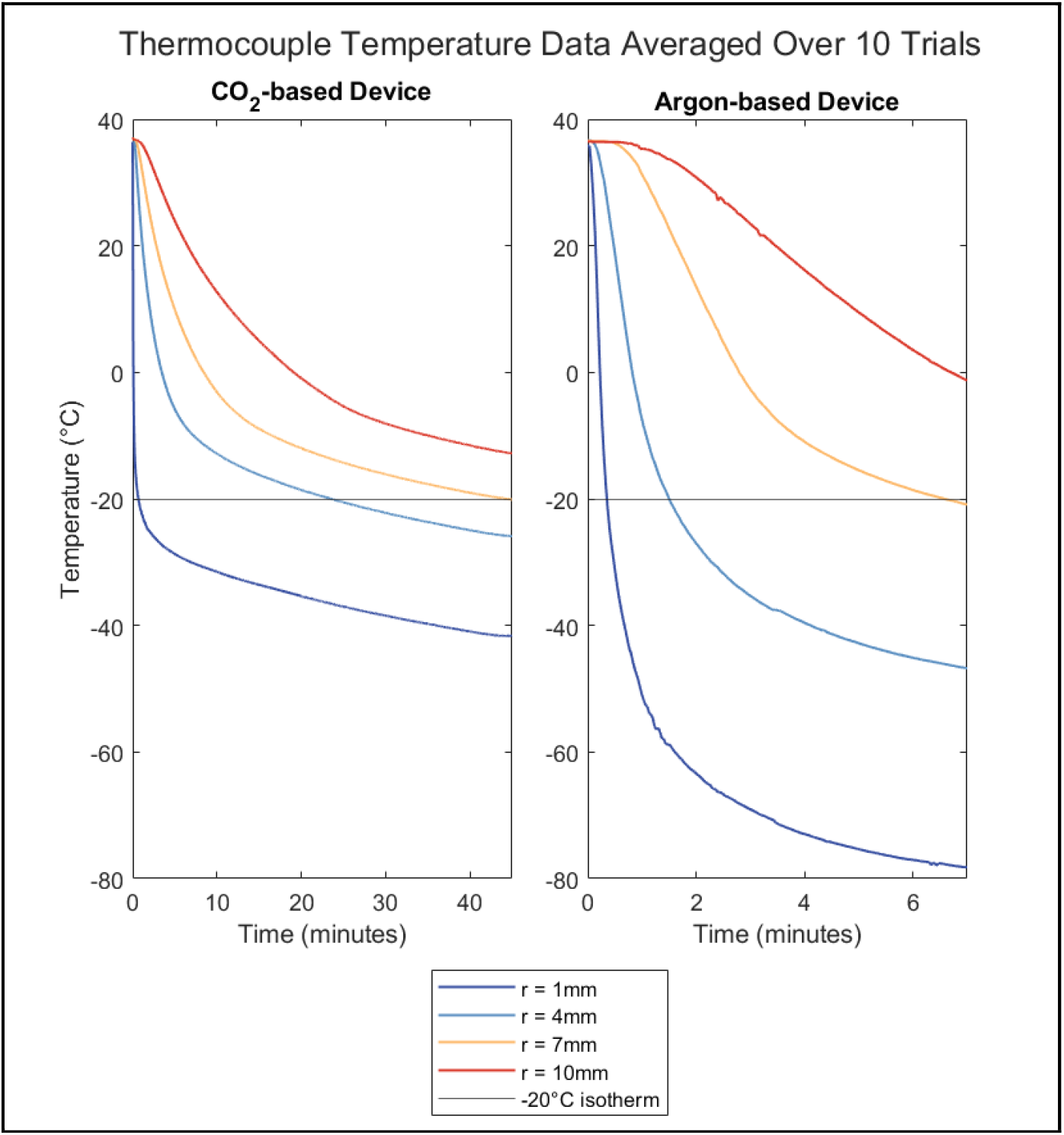
Temperature data in tissue phantom during the testing period averaged over 10 trials. *r* represents the distance of the thermocouple from the probe surface.

**Figure 6.**
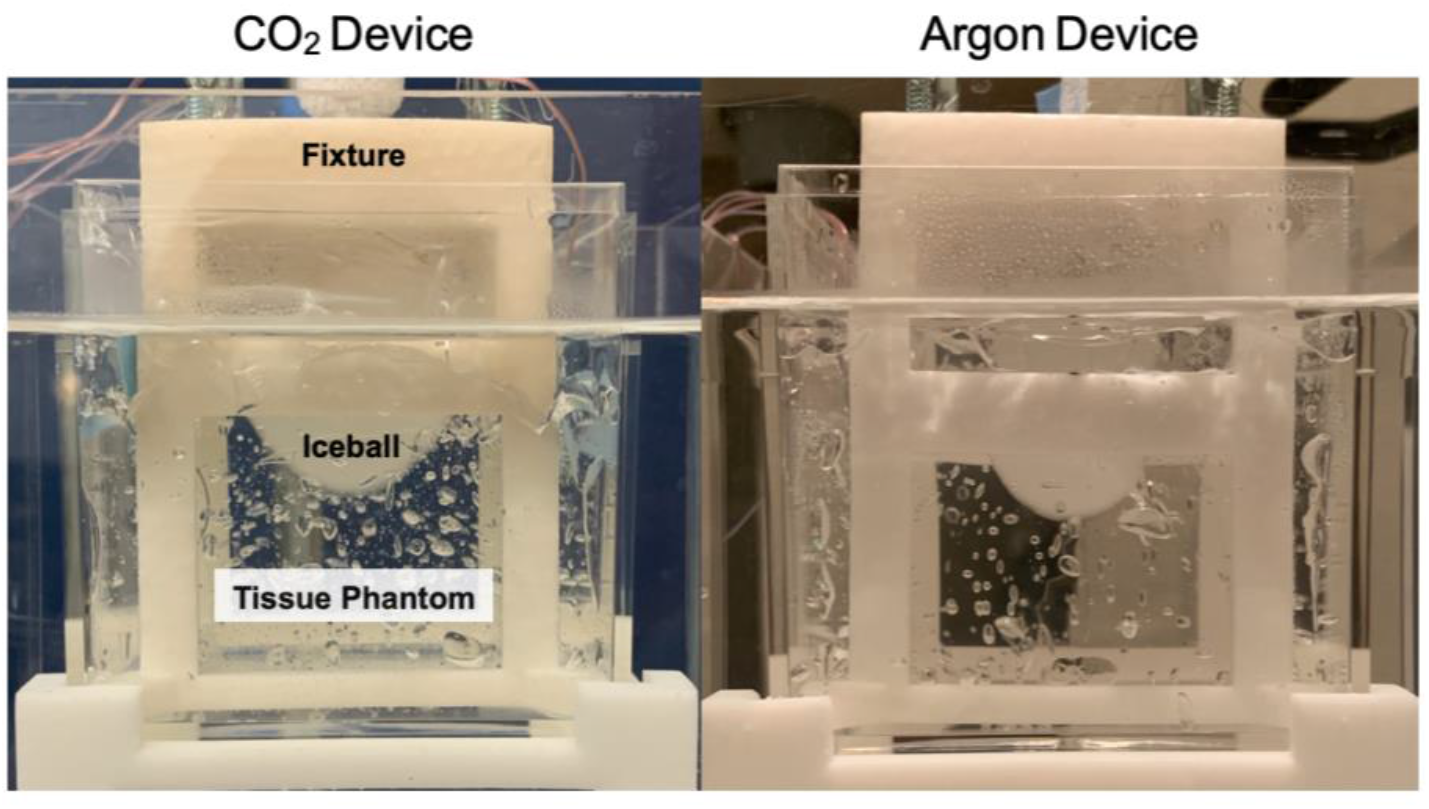
Photograph of iceballs generated during testing in the tissue phantom. Left: CO_2_ probe, right: benchmark argon probe.

The distance of the -20°C isotherm from the surface of each probe was determined using a logarithmic curve-fit (Equation 4) of the thermocouple data to represent the expected theoretical heat distribution (Figure 7). For the CO_2_ device, the distance of the - 20°C isotherm was 6.3 mm after 45 minutes, and for the argon device, the distance was 6.9 mm after 7 minutes. Accounting for the difference in probe sizes, the overall diameters of the -20°C isotherm for the CO_2_ and argon devices were 16.7 mm and 15.3 mm, respectively (Figure 8).

**Figure 7.**
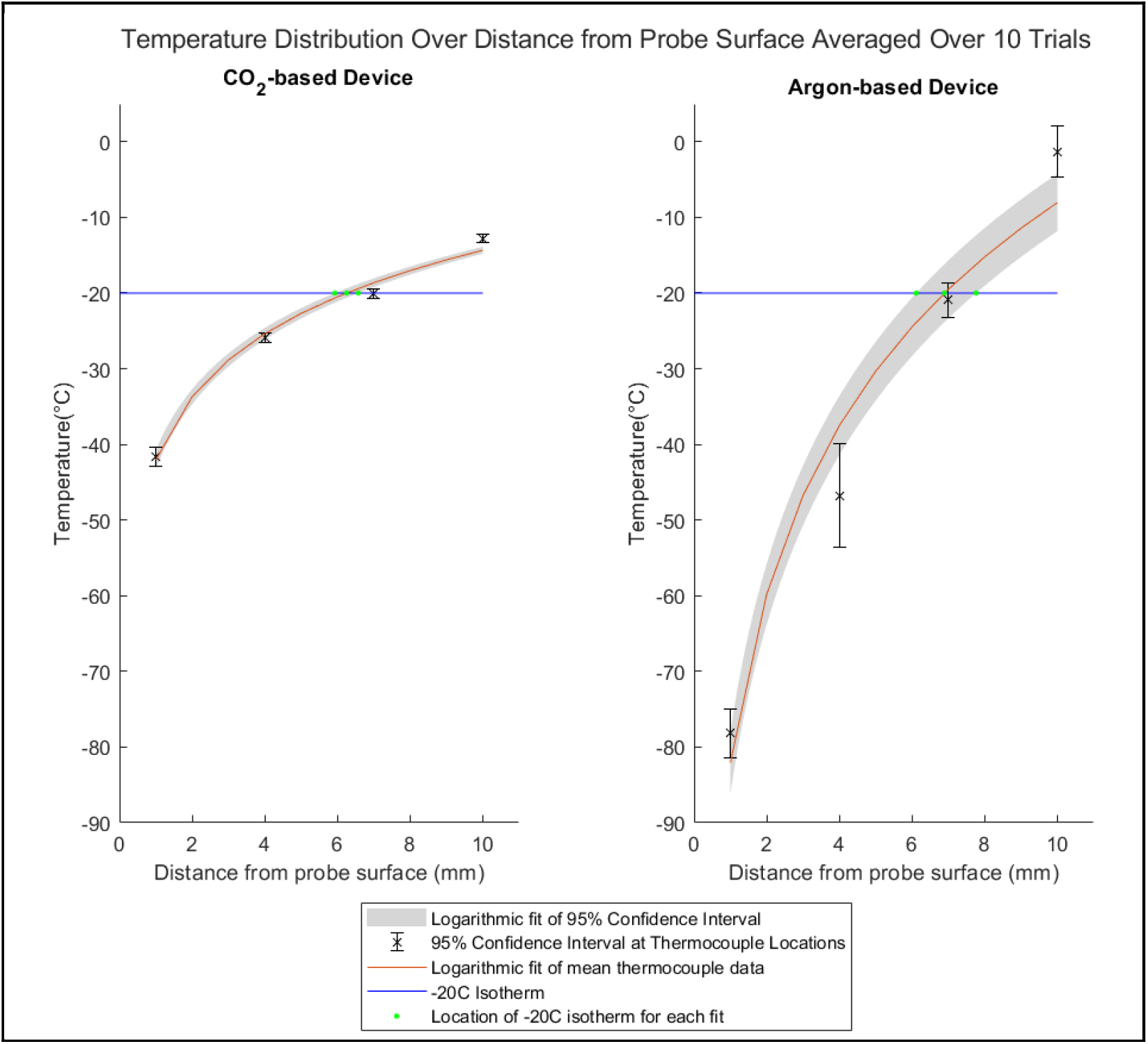
Mean and 95% confidence interval of the radial temperature distribution from the surface of the probe at the end of the testing period for each device. The red line plots the logarithmic fit of the mean thermocouple data over 10 trials. The gray area represents the 95% confidence interval of the thermocouple data fitted to a logarithmic equation. The distance of the -20°C isotherm from the surface of the probe with the 95% confidence interval is 6.3 mm (5.9, 6.6) for the CO_2_ device and 6.9 mm (6.1, 7.7) for the benchmark argon device.

**Figure 8.**
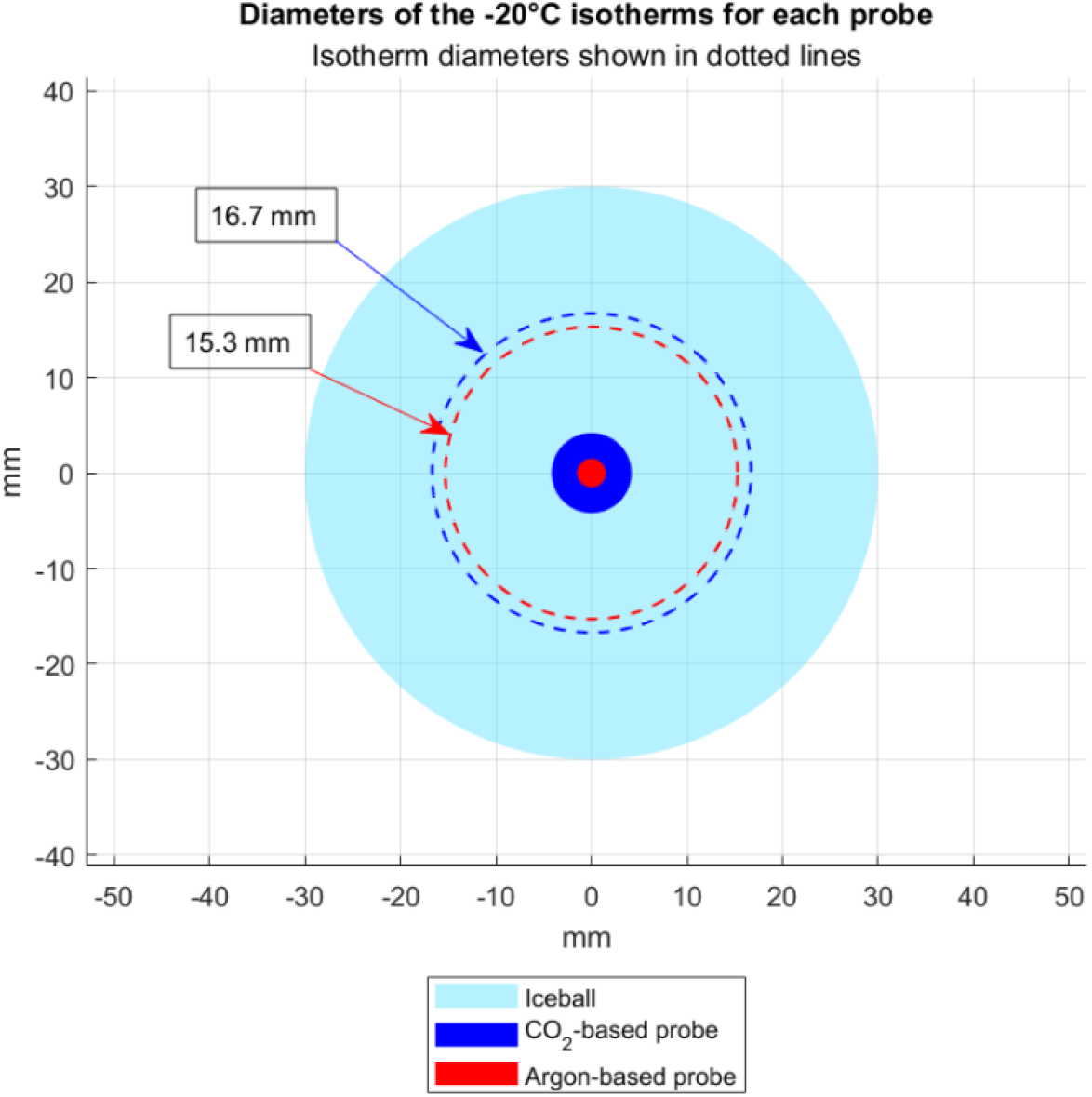
Cross-sectional representation of the probe diameters and the mean diameters of the -20°C isotherms for each probe during testing, extrapolated from the interpolated isotherm radii, assuming radial symmetry around the probe axis. Iceball size is representative.

The coefficient values *C*_*1*_ and *C*_*2*_ of the logarithmic curves (Equation 4) used to fit the average temperature data of each device were calculated with 95% confidence intervals. For the CO_2_ device, the coefficients were 12.03 [8.18, 15.88] and -42.05 [-48.44, -35.66]. For the benchmark device, the coefficients were 32.16 [11.03, 53.29] and -82.09 [-117.1, -47.03]. The goodness-of-fit of the curves was evaluated by calculating the coefficient of determination (R^2^) and the standard error (RMSE). For the CO_2_ device, R^2^ was 0.99, and the standard error was 1.57°C. For the benchmark device, R^2^ was 0.96, and the standard error was 8.61°C.

### 3.3 Histopathology Results

Figure 9 shows a photomicrograph of the H&E staining with the position of the CO_2_ cryoprobe during treatment. Because of the size limit on the pathology slides used, the tissue sample was divided into two sections for embedding and digitally reconstructed based on reference images of the fresh tissue sample immediately after resection. The pixel size of both digital images was normalized and the aspect ratio was kept constant during reconstruction (Applied Pathology Systems). Due to the tissue fixation and reconstruction process, some of the original tissue was lost along the length of the probe track. The maximum length of necrosis along the probe track, not including areas of tissue loss, was measured to be 15.01 mm. The widest diameter of necrosis perpendicular to the probe track was measured to be 15.74 mm.

**Figure 9.**
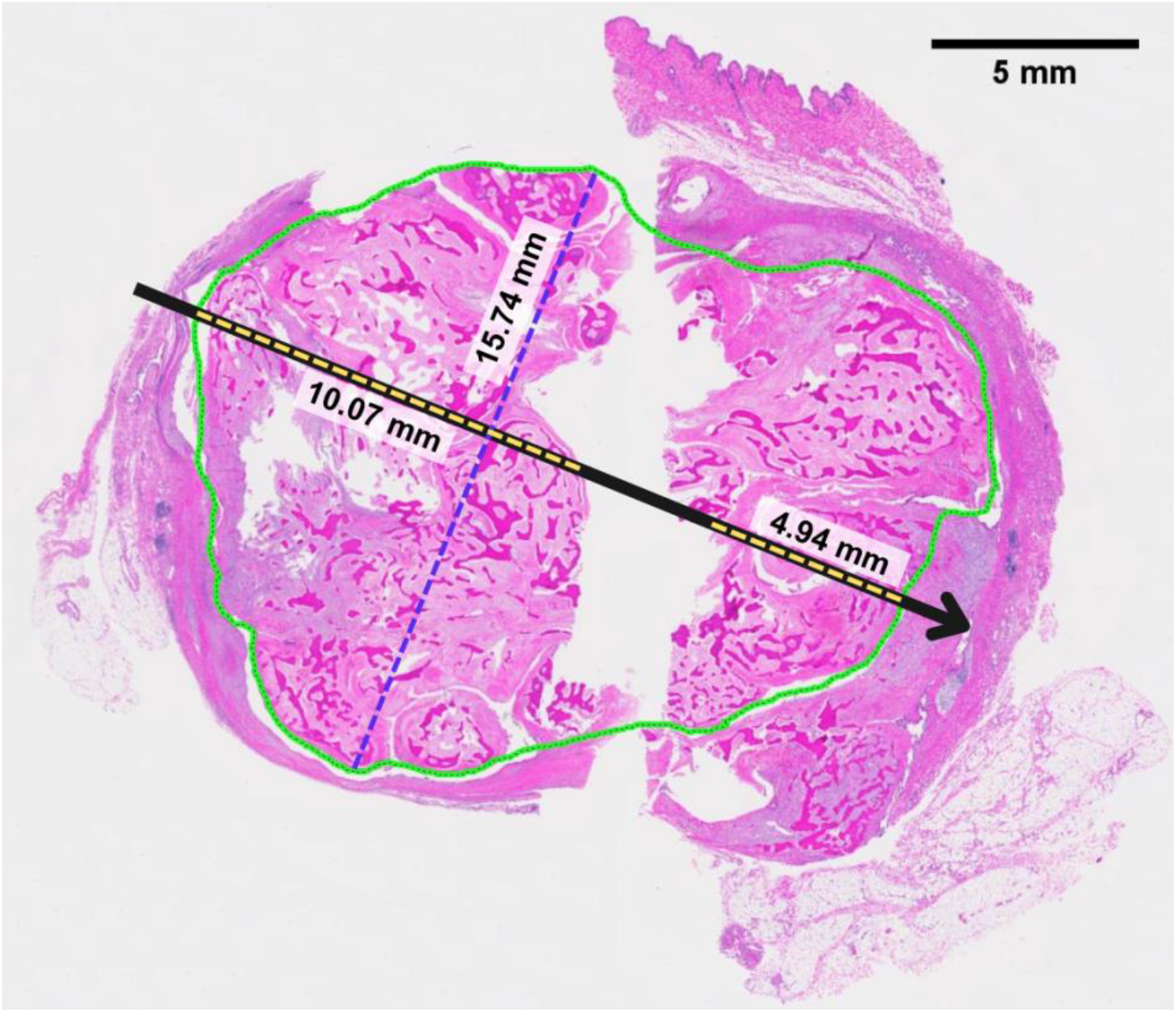
Reconstructed photomicrograph from two pieces of the mass containing annotations that show the cryoprobe orientation (black arrow). The region of necrotic tissue is outlined in green, and the maximum length and width of necrosis relative to the probe track are identified. Length measurements do not include regions without tissue between the two pieces.

## 4. Discussion

Several prior research studies correlate the size of the -20°C isotherm with the cytotoxic zone and observed margins of necrosis [23-25]. However, these studies used argon and liquid nitrogen cryotherapy devices for treatment or performed the tests in ex vivo tissue and therefore do not precisely capture the cell death mechanisms that occur during a low freeze-rate cryotherapy procedure. While temperatures below -50°C can induce cell death very quickly, holding tissue at higher temperatures of around -20°C for longer periods of time can destroy tissue through cellular dehydration and recrystallization [30-31].

The lethal temperature zone of a high freeze-rate cryotherapy device was confirmed in the argon-based device in tissue phantom testing, where a ∽1.5 cm diameter -20°C isotherm was formed in under 10 minutes. This corresponds to the indications of use for the argon-based device, which specify a lethal zone of 1.5 cm in diameter over a 10-minute treatment time [26].

Because of argon’s lower boiling point and higher working pressure as compared to CO_2_, it was anticipated that the argon device would significantly outperform the CO_2_ device when measured in a calorimeter. Although the argon device did show a marginally higher cooling power than the CO_2_ device when normalized by the active freezing surface area of the probe, the CO_2_ device demonstrated an overall cooling power of 116.9 W, a significant difference compared to the 27.7 W observed with the argon device (Table 1).

Despite the CO_2_ device having over 4 times the overall cooling power load compared to the argon device, temperature profile data indicated that the argon device established a 1.5 cm diameter -20°C isotherms much faster than the CO_2_ device (Figure 5). The difference in cooling times when compared to cooling power could be attributed to variations in thermal contact resistance between the probes as well as to the latent heat dynamics of iceball formation. Factors like contact pressure, surface area, and the thermophysical and mechanical properties of the probe materials could contribute to differences in thermal contact resistance affecting the efficiency of heat transfer [32]. Latent heat is also an important factor in iceball formation as it determines the amount of cooling energy required to form ice as the iceball grows. The latent heat of fusion for the ultrasound gel tissue phantom was estimated to be 259 J/g [33]. As the iceball radius expands, the cubic increase in ice volume demands exponentially more heat removal for continued growth. The CO_2_ device’s larger probe diameter means that its iceball has a larger initial radius than that of the argon device. This introduces a more significant latent heat barrier, requiring more cooling power for the liquid-to-solid phase change. Consequently, less cooling power is directed towards decreasing the overall temperature resulting in slower iceball growth and a longer time to generate a -20°C isotherm. These insights underscore the importance of a comprehensive evaluation of a cryotherapy probe beyond calorimetry for predicting practical temperature performance.

While the CO_2_ device took longer to generate a comparable size lethal isotherm as the benchmark device, it exhibited a significantly narrower temperature range in benchtop testing, and it also demonstrated reduced variability in temperature measurements (Table 2). A primary metric controlling the individual performance of a cryotherapy device is the pressure of the incoming gas. Because of argon’s higher cryogenic potential, the gas pressure can be downregulated during use. CO_2_, as a weaker cryogen, requires the full working pressure of a standard CO_2_ tank to achieve similar cooling outcomes. When using the CO_2_ cryotherapy device, the supplying gas tank is thermally regulated to achieve a stable starting pressure of 970 ± 30 psi, and a constant heat load is applied to the gas tank during use. This consistent approach ensures a uniform pressure drop for CO_2_ across multiple trials, albeit resulting in a gradual performance decline over time. In contrast, the direct pressure regulation of argon gas introduces irregular fluctuations in gas flow rate, potentially contributing to greater temperature variation across trials [34].

It is worth noting that this study did not explore the differences in variability between the two devices, though it presents a compelling area for future research. Additional studies may yield a deeper understanding of the factors contributing to temperature variation and their implications for diverse cryotherapy devices and applications.

Benchtop tissue phantom studies established that the CO_2_ device can generate a comparable temperature profile to the benchmark argon device. While there is a significant difference in the time for each device to reach -20°C at a 1.5 cm diameter in benchtop trials, the variability of thermal and mechanical properties between different tissue types makes it difficult to directly correlate experimental isotherm results to in vivo performance. We do not expect the actual treatment time to directly equal the time to reach a 1.5 cm diameter -20°C isotherm measured in bench testing.

Additionally, it should be noted that there are multiple factors that can cause tissue shrinkage between the time of cryotherapy treatment and the histopathological analysis of necrosis margins. After undergoing the cryotherapy procedure, excision of the treated mass was not performed until 2 weeks later. During this period, the breakdown and remodeling of necrotic tissue with fibrotic replacement is expected, leading to an overall reduction in tissue dimensions [31,35-36]. Additionally, after the mass was resected and cut into sections for analysis, the tissue sections were formalin-fixed, which can result in tissue shrinkage of 10% or more [37-38]. Due to the complex processes underlying these shrinkage factors, it is difficult to accurately quantify the level of shrinkage that occurred. As a result, it is likely that the true margin of necrosis in the body during treatment exceeds the measurement obtained through pathological analysis.

As shown in the in vivo case study, the diameter of necrosis after 20 minutes of freezing, i.e. two 10-minute freezes, with the CO_2_ device was 1.57 cm. The temperature data for the CO_2_ device show that after 20 minutes of freezing in the benchtop tissue phantom, the -20°C isotherm had a diameter of less than 1.2 cm. On the bench, the -20°C isotherm generated by the CO_2_ device took around 30 minutes to reach a 1.57 cm diameter. This suggests that the mechanisms of cell death associated with “slow” cryotherapy treatment, such as increased cellular dehydration and recrystallization, allow for a larger area of necrosis than what is predicted by bench test results. While this finding requires further validation through additional in vivo studies, it has significant implications for the development of CO_2_ cryotherapy for eventual implementation in human healthcare applications. Using CO_2_ to induce cryogenic temperatures and activate slow-cryo pathways of cell death would allow for the low-cost, effective treatment of breast cancer in low-resource regions where access to advanced treatment options is limited.

The validation of CO_2_ cryotherapy in comparison with a commercially available argon device, along with its demonstrated efficacy in necrosing clinically relevant volumes of cancerous tissue within a standard treatment time, presents a promising outlook for the application of CO_2_ cryotherapy in human treatment. Characterized by low cost and minimal resource requirements, CO_2_ cryotherapy is a compelling option for cancer treatment in low-resource settings and is well-positioned to address a growing disparity in global breast cancer care.

## Author Contributions

Conceptualization, methodology, Y.H., N.G., D.L.K., and B.S.; formal analysis, Y.H., N.G., and K.O.; investigation, Y.H., N.G., K.O., and D.L.K; resources, D.L.K, N.J.D, and B.S.; writing—original draft preparation, Y.H., N.G., and K.O.; writing—review and editing, Y.H., N.G., D.L.K., N.J.D, and B.S; project administration, B.S.; funding acquisition, Y.H. and B.S. All authors have read and agreed to the published version of the manuscript.

## Institutional Review Board Statement

The animal study protocol was approved by the Animal Care and Use Committee of Johns Hopkins University (protocol code DO21M438, date of approval 01 February 2022).

## Data Availability Statement

Data will be made available upon request.

## Acknowledgments

We would like to acknowledge Kathleen Gabrielson for her histopathology assistance, Kristen Kellar-Graney for her coordination of the study’s subject recruitment, Cheri Rice for her veterinary technician services, and Applied Pathology Systems for their pathology analysis services.

## References

1. Al-Sukhun, S.; Tbaishat, F.; Hammad, N. Breast cancer priorities in limited-resource environments: The price-efficacy dilemma in cancer care. American Society of Clinical Oncology Educational Book 2022, No. 42, 416–422 DOI: 10.1200/edbk_349861.

2. Francies, F. Z.; Hull, R.; Khanyile, R.; Dlamini, Z. Breast cancer in low-middle income countries: Abnormality in splicing and lack of targeted treatment options https://www.ncbi.nlm.nih.gov/pmc/articles/PMC7269781/ (accessed Dec 25, 2023).

3. Arnold, M.; Morgan, E.; Rumgay, H.; Mafra, A.; Singh, D.; Laversanne, M.; Vignat, J.; Gralow, J. R.; Cardoso, F.; Siesling, S.; Soerjomataram, I. Current and Future Burden of Breast Cancer: Global Statistics for 2020 and 2040. The Breast 2022, 66, 15–23. DOI:10.1016/j.breast.2022.08.010.

4. Moo, T.-A.; Sanford, R.; Dang, C.; Morrow, M. Overview of Breast Cancer Therapy. PET Clinics 2018, 13 (3), 339–354. DOI:10.1016/j.cpet.2018.02.006.

5. GlobalSurg Collaborative; National Institute for Health Research Global Health Research Unit on Global Surgery. Global variation in postoperative mortality and complications after cancer surgery: a multicentre, prospective cohort study in 82 countries. Lancet (London, England) 2021, 397 (10272), 387–397. 10.1016/S0140-6736(21)00001-5

6. Cardone, C.; Arnold, D. The Cancer Treatment Gap in Lower-to Middle-Income Countries. Oncology 2023, 101 (Suppl. 1), 2–4. DOI:10.1159/000530416.

7. Elmore, S. N.; Polo, A.; Bourque, J.-M.; Pynda, Y.; van der Merwe, D.; Grover, S.; Hopkins, K.; Zubizarreta, E.; Abdel-Wahab, M. Radiotherapy Resources in Africa: An International Atomic Energy Agency Update and Analysis of Projected Needs. The Lancet Oncology 2021, 22 (9). DOI:10.1016/s1470-2045(21)00351-x.

8. Shah, S. C.; Kayamba, V.; Peek, R. M.; Heimburger, D. Cancer Control in Low- and Middle-Income Countries: Is It Time to Consider Screening? Journal of Global Oncology 2019, No. 5, 1–8. DOI:10.1200/jgo.18.00200.

9. Donkor, A.; Atuwo-Ampoh, V. D.; Yakanu, F.; Torgbenu, E.; Ameyaw, E. K.; Kitson-Mills, D.; Vanderpuye, V.; Kyei, K. A.; Anim-Sampong, S.; Khader, O.; Khader, J. Financial Toxicity of Cancer Care in Low- and Middle-Income Countries: A Systematic Review and Meta-Analysis. Supportive Care in Cancer 2022, 30 (9), 7159–7190. DOI:10.1007/s00520-022-07044-z.

10. Newman, L. A. Breast cancer screening in low and middle-income countries. Best Practice & Research Clinical Obstetrics & Gynaecology 2022, 83, 15–23 DOI: 10.1016/j.bpobgyn.2022.03.018.

11. Ferlic, D. J.; Kotula, F. T.; Amplatz, K. Mammography apparatus, February 13, 1990.

12. Souney, S.; Roeder, R. Portable hand-carry satellite diagnostic ultrasound system for general and cardiac imaging, November 12, 2002.

13. Tran, T. T.; Hlaing, M.; Krause, M. Point-of-Care Ultrasound: Applications in Low- and Middle-Income Countries. Current Anesthesiology Reports 2021, 11 (1), 69–75. DOI:10.1007/s40140-020-00429-y.

14. Becker, D. M.; Tafoya, C. A.; Becker, S. L.; Kruger, G. H.; Tafoya, M. J.; Becker, T. K. The Use of Portable Ultrasound Devices in Low- and Middle-income Countries: A Systematic Review of the Literature. Tropical Medicine & International Health 2016, 21 (3), 294–311. DOI:10.1111/tmi.12657.

15. Bhimani, F.; Zhang, J.; Shah, L.; McEvoy, M.; Gupta, A.; Pastoriza, J.; Shihabi, A.; Feldman, S. Can the clinical utility of ibreastexam, a novel device, aid in optimizing breast cancer diagnosis? A systematic review. JCO Global Oncology 2023, No. 9 DOI: 10.1200/go.23.00149.

16. Mokbel, K.; Kodresko, A.; Ghazal, H.; Mokbel, R.; Trembley, J.; Jouhara, H. The Evolving Role of Cryosurgery in Breast Cancer Management: A Comprehensive Review. Cancers 2023, 15 (17), 4272. DOI:10.3390/cancers15174272

17. Mohammed, A.; Miller, S.; Douglas-Moore, J.; Miller, M. Cryotherapy and Its Applications in the Management of Urologic Malignancies: A Review of Its Use in Prostate and Renal Cancers. Urologic Oncology: Seminars and Original Investigations 2014, 32 (1). DOI:10.1016/j.urolonc.2013.04.004.

18. Winkle, R. A.; Mead, R. H.; Engel, G.; Kong, M. H.; Patrawala, R. A. Physician-controlled costs: The choice of equipment used for atrial fibrillation ablation. Journal of Interventional Cardiac Electrophysiology 2013, 36 (2), 157–165 DOI: 10.1007/s10840-013-9782-x.

19. Bang, H. J.; Littrup, P. J.; Goodrich, D. J.; Currier, B. P.; Aoun, H. D.; Heilbrun, L. K.; Vaishampayan, U.; Adam, B.; Goodman, A. C. Percutaneous cryoablation of metastatic renal cell carcinoma for local tumor control: Feasibility, outcomes, and estimated cost-effectiveness for palliation. Journal of Vascular and Interventional Radiology 2012, 23 (6), 770–777 DOI: 10.1016/j.jvir.2012.03.002.

20. Sprenkle, P. C.; Mirabile, G.; Durak, E.; Edelstein, A.; Gupta, M.; Hruby, G. W.; Okhunov, Z.; Landman, J. The Effect of Argon Gas Pressure on Ice Ball Size and Rate of Formation. Journal of Endourology 2010, 24 (9), 1503–1507. DOI:10.1089/end.2009.0587.

21. Cooper, S. M.; Dawber, R. P. The history of cryosurgery. Journal of the Royal Society of Medicine 2001, 94 (4), 196–201 DOI: 10.1177/014107680109400416.

22. Surtees, B.; Young, S.; Hu, Y.; Wang, G.; McChesney, E.; Kuroki, G.; Acree, P.; Thomas, S.; Blair, T.; Rastogi, S.; et al. Validation of a low-cost, carbon dioxide-based cryoablation system for percutaneous tumor ablation. PLOS ONE 2019, 14 (7) DOI: 10.1371/journal.pone.0207107.

23. Hinshaw, J. L.; Lee, F. T.; Laeseke, P. F.; Sampson, L. A.; Brace, C. Temperature isotherms during pulmonary cryoablation and their correlation with the zone of ablation. Journal of Vascular and Interventional Radiology 2010, 21 (9), 1424–1428 DOI: 10.1016/j.jvir.2010.04.029.

24. Snyder, K. K.; Van Buskirk, R. G.; Baust, J. G.; Baust, J. M. Breast cancer cryoablation: Assessment of the impact of fundamental procedural variables in an in vitro human breast cancer model. Breast Cancer: Basic and Clinical Research 2020, 14, 117822342097236 DOI: 10.1177/1178223420972363.

25. Littrup, P. J.; Freeman-Gibb, L.; Andea, A.; White, M.; Amerikia, K. C.; Bouwman, D.; Harb, T.; Sakr, W. Cryotherapy for breast fibroadenomas. Radiology 2005, 234 (1), 63–72 DOI: 10.1148/radiol.2341030931.

26. Boston Scientific Corporation. Classic Needle Sell Sheet. Boston Scientific Corporation: Marlborough, MA, n.d.; accessed 2024 Feb 26. https://www.bostonscientific.com/content/dam/bostonscientific/pi/portfolio-group/cryoablation/visualice/Classic_Needle_Sell_Sheet.pdf

27. Savitzky, Abraham.; Golay, M. J. Smoothing and differentiation of data by simplified least squares procedures. Analytical Chemistry 1964, 36 (8), 1627–1639 DOI: 10.1021/ac60214a047.

28. Hossain, S.; Mohammadi, F. One-dimensional Steady-state Analysis of Bioheat Transfer Equation: Tumour Parameters Assessment for Medical Diagnosis Application. IMETI 2013 - 6th International Multi-Conference on Engineering and Technological Innovation, Proceedings 2013, 26–30.

29. Parsons, K.; Reichanadter, T.; Vicksman, A.; Segur, H. Explicit solution for cylindrical heat conduction. Volume 13, Issue 2 2016, 13 (2) DOI: 10.33697/ajur.2016.020.

30. Theodorescu D. Cancer cryotherapy: evolution and biology. Rev Urol. 2004;6(Suppl 4):S9–S19.

31. Mazur P. Physical-chemical factors underlying cell injury in cryosurgical freezing. In: Rand R, Rinfret A, von Leden H, editors. Cryosurgery. Charles C. Thomas; Springfield, IL.: 1968. pp. 32–51.

32. Salmon, K.; Nellis, G.; Pfotenhauer, J. Experimental characterization of cryogenic contact resistance. Cryogenics 2022, 128, 103587 DOI: 10.1016/j.cryogenics.2022.103587.

33. Etheridge, M. L.; Choi, J.; Ramadhyani, S.; Bischof, J. C. Methods for characterizing convective cryoprobe heat transfer in Ultrasound Gel Phantoms. Journal of Biomechanical Engineering 2013, 135 (2) DOI: 10.1115/1.4023237.

34. Understanding the characteristics of pressure regulators https://fluidpowerjournal.com/understanding-the-characteristics-of-pressure-regulators/ (accessed Dec 28, 2023).

35. Erinjeri, J. P.; Clark, T. W. I. Cryoablation: Mechanism of action and Devices. Journal of Vascular and Interventional Radiology 2010, 21 (8) DOI: 10.1016/j.jvir.2009.12.403.

36. Kwak, K.; Yu, B.; Lewandowski, R. J.; Kim, D.-H. Recent progress in cryoablation cancer therapy and nanoparticles mediated cryoablation. Theranostics 2022, 12 (5), 2175–2204 DOI: 10.7150/thno.67530.

37. Tran, T.; Sundaram, C. P.; Bahler, C. D.; Eble, J. N.; Grignon, D. J.; Monn, M. F.; Simper, N. B.; Cheng, L. Correcting the shrinkage effects of formalin fixation and tissue processing for renal tumors: Toward standardization of pathological reporting of Tumor Size. Journal of Cancer 2015, 6 (8), 759–766 DOI: 10.7150/jca.12094.

38. Upchurch, D. A.; Klocke, E. E.; Henningson, J. N. Amount of skin shrinkage affecting tumor versus grossly normal marginal skin of dogs for cutaneous mast cell tumors excised with curative intent. American Journal of Veterinary Research 2018, 79 (7), 779–786 DOI: 10.2460/ajvr.79.7.779.

